# Protein proximity networks and functional evaluation of the Casein Kinase 1 *γ* family reveals unique roles for CK1*γ*3 in WNT signaling

**DOI:** 10.1101/2021.11.30.470617

**Authors:** Megan J. Agajanian, Frances M. Potjewyd, Brittany M. Bowman, Smaranda Solomon, Kyle M. LaPak, Dhaval P. Bhatt, Jeffrey L. Smith, Dennis Goldfarb, Alison D. Axtman, Michael B. Major

## Abstract

The WNT/*β*-catenin signaling pathway is evolutionarily conserved and controls normal embryonic development, adult tissue homeostasis and regeneration. Aberrant activation or suppression of WNT signaling contributes to cancer initiation and progression, developmental disorders, neurodegeneration, and bone disease. Despite great need and more than 40 years of research, targeted therapies for the WNT pathway have yet to be fully realized. Kinases are exceptionally druggable and occupy key nodes within the WNT signaling network, but several pathway-relevant kinases remain understudied and ‘dark’. Here we studied the function of the CSNK1*γ* subfamily of human kinases. miniTurbo-based proximity biotinylation and mass spectrometry analysis of CSNK1*γ*1, CSNK1*γ*2, and CSNK1*γ*3 revealed numerous established components of the *β*-catenin- dependent and independent WNT signaling pathway, as well as novel interactors. In gain-of-function experiments leveraging a panel of transcriptional reporters, CSNK1*γ*3 but not CSNK1*γ*1 or CSNK1*γ*2 activated *β*-catenin-dependent WNT signaling and the Notch pathway. Within the family, CSNK1*γ*3 expression uniquely induced LRP6 phosphorylation. Conversely, siRNA-mediated silencing of CSNK1*γ*3 alone had no impact on WNT signaling, though co-silencing of all three family members decreased WNT pathway activity. We characterized two moderately selective and potent small molecule inhibitors of the CSNK1*γ* family. These inhibitors and a CSNK1*γ*3 kinase dead mutant suppressed but did not eliminate WNT-driven LRP6 phosphorylation and *β*-catenin stabilization. Our data suggest that while CSNK1*γ*3 expression uniquely drives pathway activity, potential functional redundancy within the family necessitates loss of all three family members to suppress the WNT signaling pathway.

## Introduction

WNT signaling regulates embryonic development, injury repair, regeneration, and tissue homeostasis (1, 2). WNT signaling mechanisms are tightly controlled by feedback loops and rapid protein degradation. In disease states including cancer, neurodegeneration and bone density disorders, regulatory mechanisms governing WNT signaling are aberrantly active or inhibited (3–6). In some cancers mutations in core signaling proteins are common, including APC mutations in greater than 80% of colorectal adenocarcinoma. In other cancers of the lung and breast, increased WNT ligand secretion from stromal cells drives pathway activity (5,7–9). While WNT signaling has been well-studied across many disease states and several therapeutics have been developed, targeting WNT clinically remains difficult (1,4,10,11). Identifying and understanding new regulators of WNT signaling may reveal therapeutic targets to improve patient outcome for WNT-associated diseases.

Kinases, a highly druggable class of proteins, are master regulators of various signaling pathways, including WNT signaling. WNT/*β*-catenin dependent signaling is often described with respect to the main effector protein, *β*-catenin. In the absence of WNT ligand, a destruction complex, comprised of CK1*α* (CSNK1A1), GSK3*β*, AXIN, and APC, phosphorylates *β*-catenin resulting in its binding to and subsequent ubiquitylation and degradation by *β*-TRCP and the proteasome (12–17). In the presence of WNT ligand, the WNT receptors LRP6 and Frizzled co-complex and an alternate signaling complex, the WNT signalosome, forms around the intracellular tail of LRP6. Signalosome formation transiently suppresses *β*-catenin phosphorylation, allowing *β*-catenin to accumulate, translocate to the nucleus and drive transcription of WNT response genes (18–23). Several kinases (CSNK1 family, GSK3*β*, RIPK4, AAK1, the GRK family) regulate WNT signaling in both the signalosome and the destruction complex (24–29). Of these kinases, GSK3*β* has been the most thoroughly studied within the WNT field and as a master regulator of cell signaling networks (24, 30). The CSNK1 family (referred to as CK1 hereafter), while also widely studied, is more complex. The CK1 family is comprised of CK1*α* (CSNK1A), CK1*δ* (CSNK1D) CK1*ε* (CSNK1E), and CK1*γ* (CSNK1G). CK1*α* functions almost exclusively within the destruction complex where it phosphorylates *β*-catenin to prime for phosphorylation by GSK3*β* (17). This triggers the ubiquitylation and degradation of *β*-catenin. CK1*δ* and CK1*ε* function redundantly within the destruction complex and within the WNT signalosome (31, 32). The CK1*γ* subfamily, as well as CK1*δ* and CK1*ε*, have a reported role in priming LRP6 for GSK3*β* phosphorylation and subsequent signalosome formation (31, 32).

The CK1*γ* subfamily of kinases is comprised of three genes: CK1*γ*1 (CSNK1G1), CK1*γ*2 (CSNK1G2), and CK1*γ*3 (CSNK1G3). All three are identified as understudied and dark kinases by the NIH initiative, Illuminating the Druggable Genome (IDG; https://commonfund.nih.gov/idg/index) (33). The CK1*γ* subfamily share 92% sequence homology within their kinase domain. The homology in their C-terminal regulatory domain is 41% conserved, with CK1*γ*3 containing a unique 33 amino acid insertion (**Fig. 1A**). Studies suggest that the palmitoylation modification positions them to act within the WNT signalosome, as other CK1 family members are not anchored to the membrane (31). Apart from an emerging role of CK1*γ* in WNT/*β*-catenin signaling, the CK1*γ* family has described roles in promoting phosphorylation of p65, (NF-*κ*B subunit), resulting in inhibition of the retinoic acid-inducible gene 1 (RIG-1) mediated signaling (34). Additionally, CK1*γ* can activate TNF*α* signaling (35). In *Drosophila*, Giglamesh (CK1*γ Drosophila* homolog) has an established role in *β*-catenin independent signaling in regulating Van Gogh phosphorylation at the membrane as well as regulating NOTCH pathway activity (36, 37).

**Figure 1.**
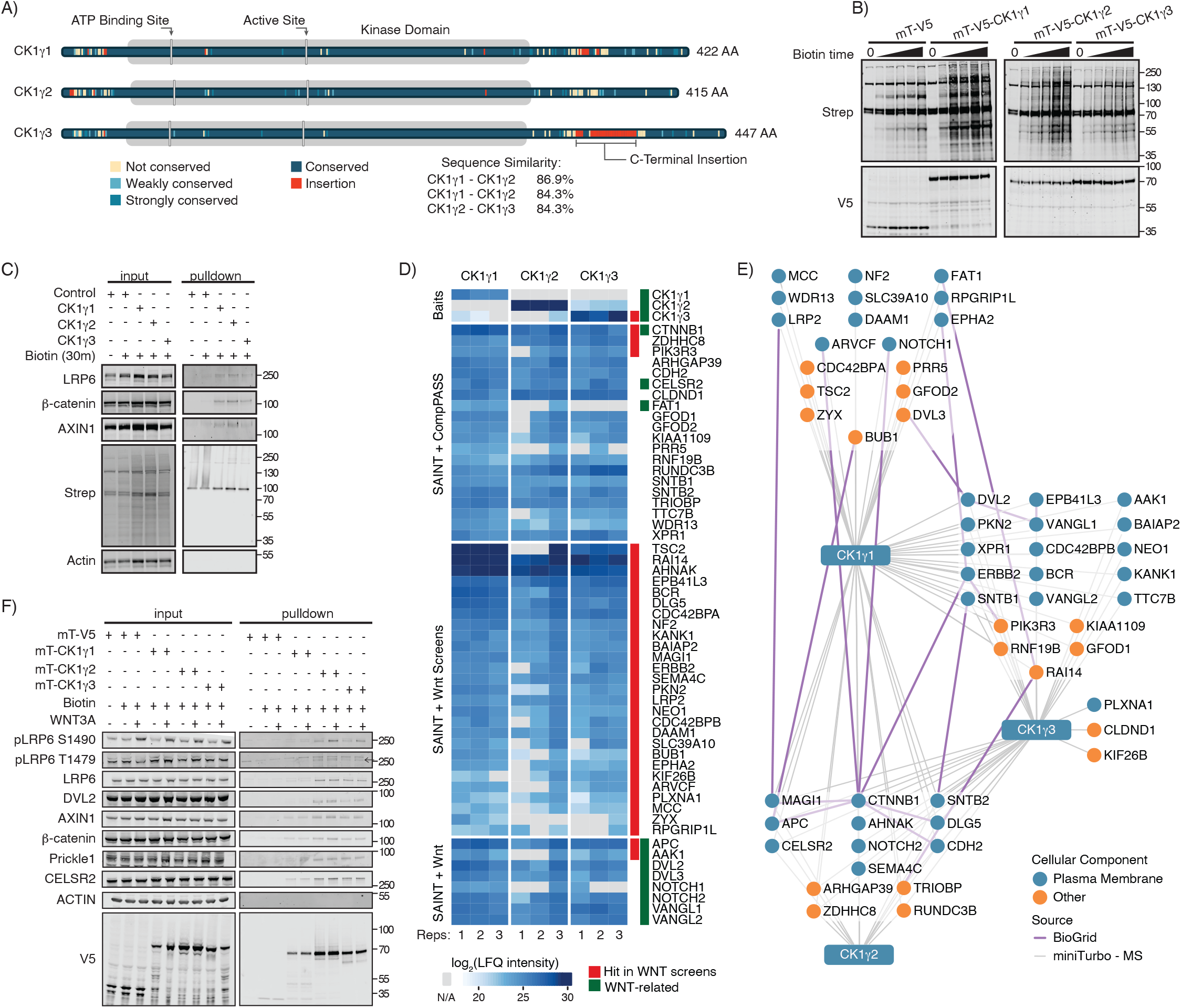
CK1γ proximity-based interaction networks identify WNT components. **(A)** Schematic of CK1γ family highlighting conserved amino acids, sequence similarity between family members and mutations. **(B)** W. blot analysis of stable HEK293T cells expressing mini-Turbo CK1γ or V5 control incubated with 50 μM of biotin over a time course (^α^untreated, 15 min, 30 min, 45 min, 1 hr, 2 hr). **(C)** W. blot analysis of streptavidin affinity pulldown from stable HEK293T cells expressing mini-Turbo CK1γ or V5, treated as indicated. **(D)** Heat map demonstrating changes in log2(LFQ intensity) of protein interactions of CK1γ subfamily AP-MS. Red track indicates hits in previously published WNT screens **(Table S1, S2)**. Green track indicates proteins involved in WNT/β-catenin-dependent and independent signaling. **(E)** Affinity purification mass spectrometry protein-protein interaction networks for CK1γ1, CK1γ2, CK1γ. HEK293T cells stably expressing mT-V5-CK1γ construct were treated with 50 μM biotin for 30 mins. Blue circle indicates localization to the plasma membrane, and orange circles represent alternate subcellular localizations. Grey lines represent bait-prey interactions, while purple lines indicate prey-prey interactions determined by BioGrid. (F) W. blot validation of mT-CK1γ protein-protein interaction networks treated with biotin and WNT3A CM for 30 mins.

Following WNT ligand engagement of FZD, DVL is recruited to the membrane and binds FZD (38). In turn, this promotes recruitment of the AXIN complex (GSK3*β*, CK1 and APC) to LRP6, forming the WNT signalosome (21,39,40). Central to WNT signalosome formation is phosphorylation of LRP6 at multiple sites, with GSK3*β* phosphorylating LRP6 at S1490 and CK1 phosphorylating LRP6 at T1479 and T1493 (31, 32). Ultimately, CK1 phosphorylation of LRP6 triggers AXIN recruitment (31,32,41,42). Upon signalosome formation, GSK3*β* is sterically occluded by the cytoplasmic tail of phosphorylated LRP6, transiently suppressing *β*-catenin phosphorylation (21,23,31,32). Though several studies have identified the CK1 family as essential regulators of LRP6 phosphorylation and establishment of the signalosome, important questions remain. For example, each member of the CK1*γ* subfamily has not been individually evaluated within *β*-catenin-dependent WNT signaling. Additionally, previously reported data relied exclusively on overexpression studies and presented no genetic or chemical loss-of-function data (31, 32).

Here we defined protein-protein proximity networks for the CK1*γ* family and functionally evaluated their contribution to a small panel of disparate signaling pathways. Each CK1*γ* family member co-complexed with various WNT components, including *β*-catenin and core members of the planar cell polarity complex. Surprisingly, only CK1*γ*3 activated *β*-catenin-dependent WNT signaling when overexpressed. Our experiments reveal a family-specific role for CK1*γ*3 in promoting phosphorylation of LRP6 within the WNT signalosome. Additionally, we characterized two pan-CK1*γ* specific inhibitors *in vitro* and in cell-based assays. Inhibition of CK1*γ* impairs the ability of WNT signaling to be fully activated by WNT ligand. This finding was extended and confirmed with a kinase dead CK1*γ*3 mutant, demonstrating that CK1*γ*3 kinase activity is required to activate WNT signaling. Overall, this work establishes that CK1*γ*3 positively regulates WNT signaling.

## Results

### Physical and functional evaluation of the CK1γ family

To better understand each member of the CK1*γ* family, we defined their protein-protein proximity networks and functionally evaluated their impact on signal transduction. We used the miniTurbo (mT) promiscuous biotin ligase to comprehensively map CK1*γ* proximal proteins, which requires a shorter biotin labeling window compared to BirA* (43, 44). Specifically, in the presence of exogenous biotin, a protein of interest fused to the mT biotin ligase will result in biotinylation of surface exposed lysine residues on proximal proteins, enabling their affinity purification with streptavidin and identification by Western blot (W. blot) or mass spectrometry (MS). HEK293T stably expressing mT-CK1*γ*1, mT-CK1*γ*2, mT-CK1*γ*3 were treated with biotin for varying amounts of time before W. blot analysis for biotinylated proteins and for a V5 epitope, the latter of which is cloned in frame with mT (**Fig. 1B**). Streptavidin affinity purification followed by W. blot analysis confirmed proximal complex formation between CK1*γ* family members and WNT pathway proteins LRP6, AXIN1 and *β*-catenin (CTNNB1) (**Fig. 1C**). We next used MS across biological triplicate experiments to identify and quantify proximal proteins to each of the CK1*γ* family members. Resulting identifications were probabilistically scored against controls with SAINT and then further ranked with CompPASS before visualization (**Table S1, sheet B, C**) (45–47). Heat map representation of protein abundance revealed concordant proximal networks across the CK1*γ* family (**Fig. 1D**). Of 545 high confidence protein interactions for CK1*γ*1, CK1*γ*2, CK1*γ*3, 326 proteins were seen with all subfamily members. *β*-catenin, ZDHHC8 (Zinc Finger DHHC-Type Palmitoyltransferase 8), and PIK3R3 (Phosphoinositide-3-Kinase Regulatory Subunit 3) were the most abundant proximal proteins in all three networks. Several core components of the *β*-catenin dependent and independent signaling pathway were also identified, such as DVL, APC, CELSR2, and VANGL (**Fig. 1D, green track**). To further explore these networks and their connectivity to the WNT pathway, we annotated all proteins for functional contribution to *β*-catenin-dependent transcription. From four recently published independent genetic screens, we identified a subset of genes as functionally impactful to *β*-catenin-dependent transcription (**Table S2**) (48–51). Integration of these data with the CK1*γ* proximity networks revealed 32 CK1*γ* proximal proteins that when silenced or CRISPR-deleted impacted *β*-catenin- dependent transcription. (**Fig. 1D, red track**). Last, we visualized the data as a network to examine high confidence prey-prey interactions and subcellular localization (**Fig. 1E**). The majority of the prey proteins are known to localize to the plasma membrane.

To confirm the MS results, and to test the impact of WNT3A stimulation, we analyzed the CK1*γ* affinity purified proximal proteins by W. blot analysis (**Fig. 1F**). CK1*γ*2 and CK1*γ*3 robustly interacted with *β*-catenin-dependent WNT signaling components (LRP6, DVL2, AXIN1 and *β*- catenin), while CK1*γ*1 interacted with these proteins but to a lesser extent. WNT3A stimulation increased pLRP6 S1490 in all three CK1*γ* pulldowns. Additional WNT3A-dependent changes were not observed. We also confirmed the interaction between the CK1*γ* family and proteins involved in *β*-catenin independent signaling, CELSR2 and PRICKLE1 (**Fig. 1F**).

We next functionally evaluated each member of the CK1*γ* subfamily across a small panel of engineered pathway-specific transcriptional reports. Transient transfection and over-expression of each CK1*γ* subfamily member in HEK293T cells revealed relationships to the WNT (BAR), NOTCH, TGF*β* (SMAD), NRF2 (hQR41), and RA (retinoic acid, RAR), and TNF*α* (NF*κ*B) signaling pathways (**Fig. 2A**) (27,52–54). Surprisingly, only CK1*γ*3 activated the WNT reporter, with minimal effects from CK1*γ*1 and CK1*γ*2. Though statistically significant compared to the negative control, none of the CK1*γ* family members strongly regulated the other signal transduction reporters. To further establish the CK1*γ*3 selectivity for *β*-catenin-dependent activation of the BAR reporter, we performed a dose response overexpression experiment. In contrast to CK1*γ*3, overexpression of CK1*γ*1 and CK1*γ*2 did not activate BAR (**Fig. 2B, C**). Similar to previously published reports, in the presence of WNT3A conditioned media (CM), CK1*γ*1 and CK1*γ*2 activated WNT signaling, albeit not to the same extent as CK1*γ*3 (**Fig. 2D**) (31, 32). Additionally, co-overexpression of LRP6 with either CK1*γ*1, CK1*γ*2, or CK1*γ*3 resulted in a significant activation of BAR compared to control and compared to LRP6 overexpression alone (**Fig. 2E**) (31, 32). Together, these data establish CK1*γ*3 as proximal to numerous WNT pathway proteins and as an activator of *β*-catenin-dependent WNT signaling. CK1*γ*1 and CK1*γ*2 share WNT pathway proximal proteins, but in the absence of WNT3A or co-overexpression of LRP6, they do not regulate *β*-catenin-dependent transcription.

**Figure 2.**
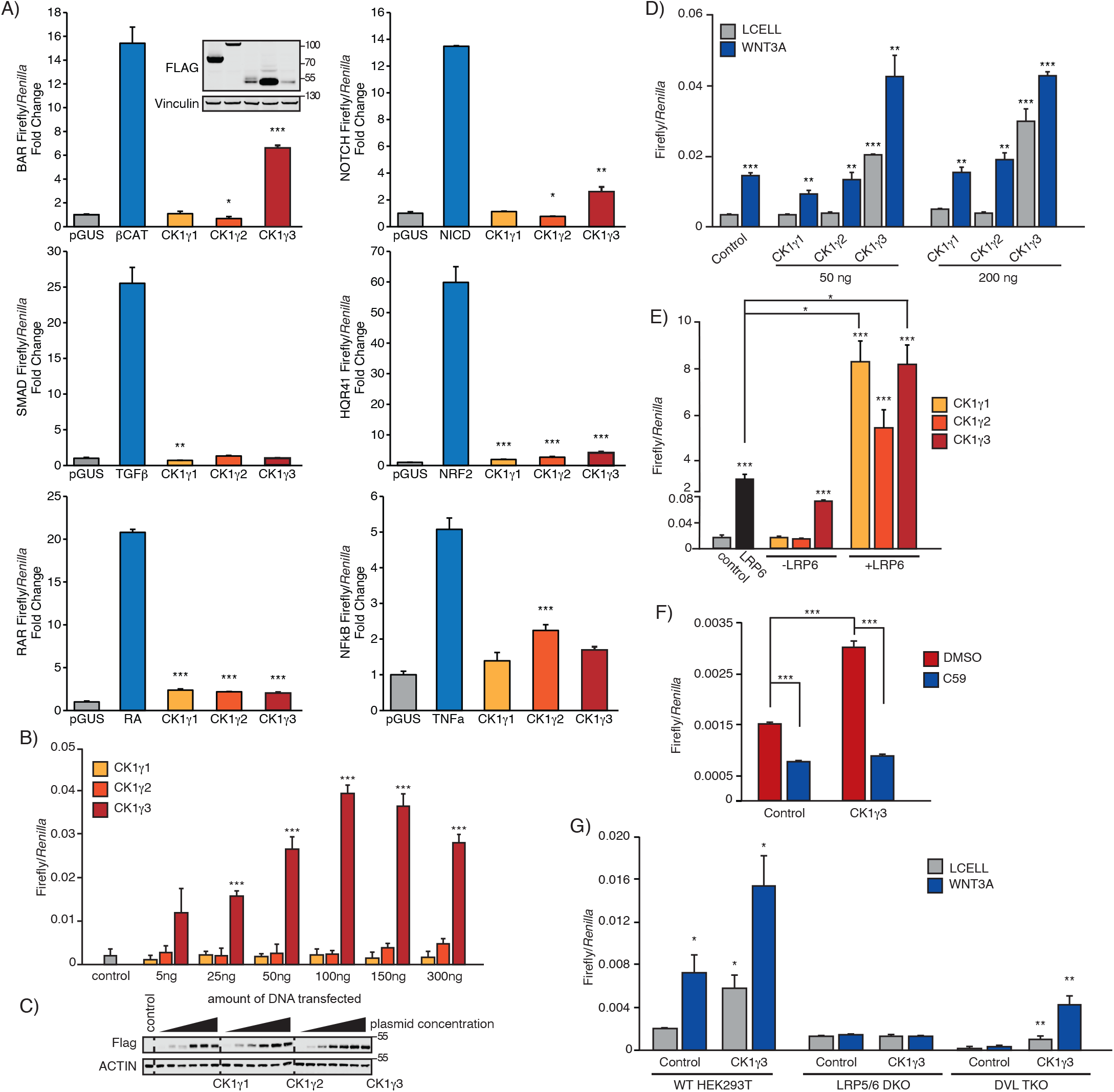
Low throughput reporter screen identifies CK1γ3 as an activator of WNT signaling. **(A)** HEK293T cells transfected for 30 hrs with 100 ng of indicated control (β-catenin, NRF2, NCID) construct or pGUS control or indicated CK1γ construct then 40 ng TK-*Renilla* and 40 ng Response Element-Luciferase. For treatments (TGFβ 10 ng/mL, TNFα 10 ng/mL, RA), cells were treated for 6 hrs (TGFβ, TNFα) or 16 hrs (RA). Inset W. blot demonstrates overexpression for BAR luciferase assay. Each condition was normalized to pGUS. Error bars represent standard deviation and statistics are compared to pGUS control. **(B)** BAR luciferase assay from HEK293T B/R (stable BAR-firefly luciferase, TK-*Renilla* expressing cell line) cells transfected for 24 hrs with indicated doses of construct. Statistics are compared to pGUS control. **(C)** Confirmation of overexpression for luciferase assay in panel B. **(D)** HEK293T B/R cells were transfected with the indicated constructed for 14 hrs, then treated with Lcell or WNT3A CM for 16 hrs. All statistics are compared to Flag-Control Lcell sample. **(E)** HEK293T B/R cells were transfected with the indicated constructed for 24 hrs then analyzed by luciferase assay. Statistics are compared to pGUS control. **(F)** HEK293T B/R cells were transfected with the indicated constructed for 14 hrs, then treated with 10 μM C59 for 18 hrs. **(G)** Wildtype, LRP5/6 DKO, DVL TKO HEK293T cells transfected with 20 ng BAR-luciferase, 10 ng TK-*Renilla* and 70 ng of indicated constructs for 14 hrs, then treated with either Lcell CM or WNT3A CM for 18 hrs. Statical significance presented are compared to Lcell treated Flag-Control cells for each cell type. All panels *** p<0.0005, ** p<0.005, * p<0.05 and all data are representative of biological triplicates, unless otherwise noted. Error bars represent S.E. Bars represent average Firefly/*Ren* (RFU) from 3 technical replicates +/- standard error (S.E.), unless otherwise stated.

### CK1γ3 requires WNT components to activate WNT signaling

To evaluate which WNT pathway proteins are required for CK1*γ*3 activation of *β*-catenin- dependent transcription, we studied established inhibitors of WNT signaling and knockout cell lines. While overexpression of CK1*γ*3 activated BAR in the absence of WNT ligand, treatment with C59, a PORCN inhibitor (required for WNT ligand palmitoylation and secretion), blocked this activation, which supports a role for autocrine WNT signaling in HEK293T cells (**Fig. 2F**) (55). We next tested the requirement of LRP6 and DVL. In contrast to HEK293T wildtype cells, overexpression of CK1*γ*3 in LRP5/6 double knockout (DKO) HEK293T cells did not activate WNT signaling (**Fig. 2G**). HEK293T cells lacking DVL1, DVL2 and DVL3 did not respond to WNT3A CM, as expected (56). Surprisingly, CK1*γ*3 overexpression increased BAR activity in DVL KO cells, but to a lesser extent that in DVL wild type cells (**Fig. 2G**). These data indicate that WNT3A ligand secretion and expression of LRP5/6 are required for CK1*γ*3-driven activation of *β*-catenin-dependent transcription; the DVL proteins are involved but are not absolutely required.

### CK1γ3 activates WNT signaling by increasing LRP6 phosphorylation

It was previously reported that overexpression of CK1*γ*1 together with overexpression of LRP6 and DVL resulted in LRP6 phosphorylation at T1493 and T1479 (31,32,57). In HEK293T cells, we tested whether overexpression of each member CK1*γ* family impacted LRP6 phosphorylation (**Fig. 3A**). CK1*γ*1 overexpression did not increase LRP6 phosphorylation at T1479 or the GSK3*β* site, S1490, irrespective of WNT3A treatment (**Fig. 3B, C**). With CK1*γ*2 overexpression, S1490 LRP6 phosphorylation increased in response to WNT3A, but not T1479. CK1*γ*3 overexpression robustly increased LRP6 phosphorylation at T1479, as well as S1490, in both the absence and presence of WNT3A. Together these data suggest that in contrast to CK1*γ*3, CK1*γ*1 and CK1*γ*2 do not activate WNT signaling in the absence of exogenous WNT3A ligand and do not induce LRP6 phosphorylation at T1479.

**Figure 3.**
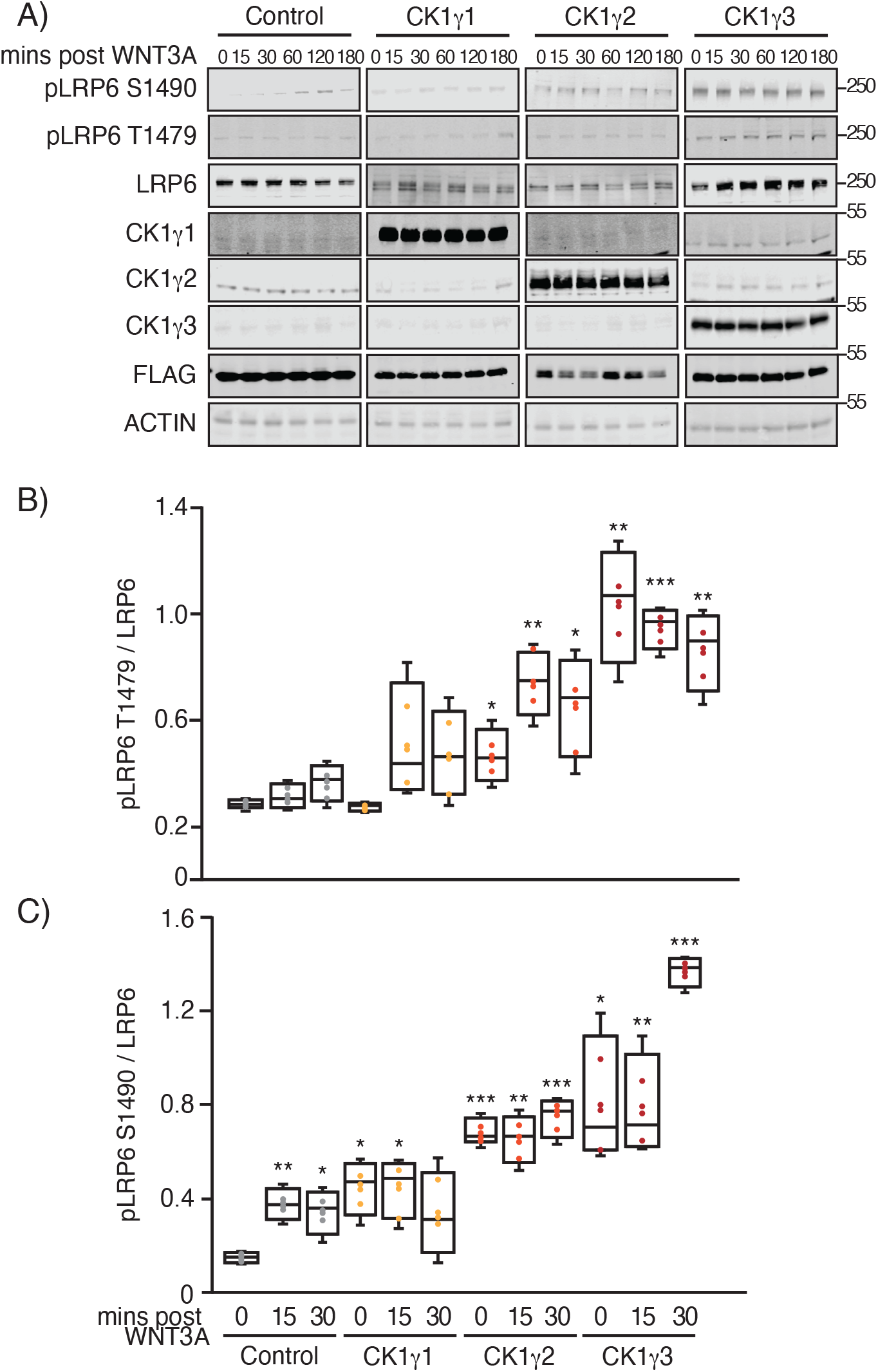
Overexpression of CK1γ3 activates WNT signaling by increasing LRP6 phosphorylation. **(A)** HEK293T cells were transfected with the indicated constructed for 14 hrs, then exposed to WNT3A CM for indicated time and analyzed by W. blot. **(B, C)** Box-and-whisker plot quantification of W. blot from panel A for indicated time points, pLRP6 T1479 (panel B) and pLRP6 S1490 (panel C) across 4 replicates, normalized to total LRP6. For all panels: *** p<0.0005, ** p<0.005, * p<0.05 and all data are representative of biological quadruplicates. All statistics are compared to Flag-Control untreated sample.

### CK1γ3 kinase activity was required for maximum activation of WNT signaling

We next performed loss-of-function studies to determine if the CK1*γ* family was required for WNT3A-driven *β*-catenin-dependent transcription. First, we generated a CK1*γ*3 kinase dead mutant (CK1*γ*3-K72A). In contrast to CK1*γ*3-WT, expression of kinase-dead CK1*γ*3-K72A in HEK293T cells did not regulate the activity of the BAR transcriptional reporter (**Fig. 4A**). In the presence of WNT3A CM, kinase dead CK1*γ*3-K72A expression suppressed BAR activity as compared to wild type CK1*γ*3 or control (**Fig. 4A**). In an orthogonal experiment, CK1*γ*3 or CK1*γ*3- K72A were expressed in HEK293T cells stably harboring a BAR-GFP fluorescent reporter. Live cell imaging over three days revealed that CK1*γ*3-K72A suppressed WNT3A-driven *β*-catenin- dependent transcription (**Fig. 4B**). Last, we tested whether expression of the CK1*γ*3 kinase dead mutant impacted the phosphorylation of LRP6. CK1*γ*3 overexpression resulted in LRP6 phosphorylation at both S1490 and T1479 sites following WNT3A CM exposure (**Fig. 4C-E**). In contrast, overexpression of CK1*γ*3-K72A suppressed phosphorylation of LRP6 at both sites. These data demonstrate that kinase activity of CK1*γ*3 is required for phosphorylation of LRP6 and activation of the WNT pathway.

**Figure 4.**
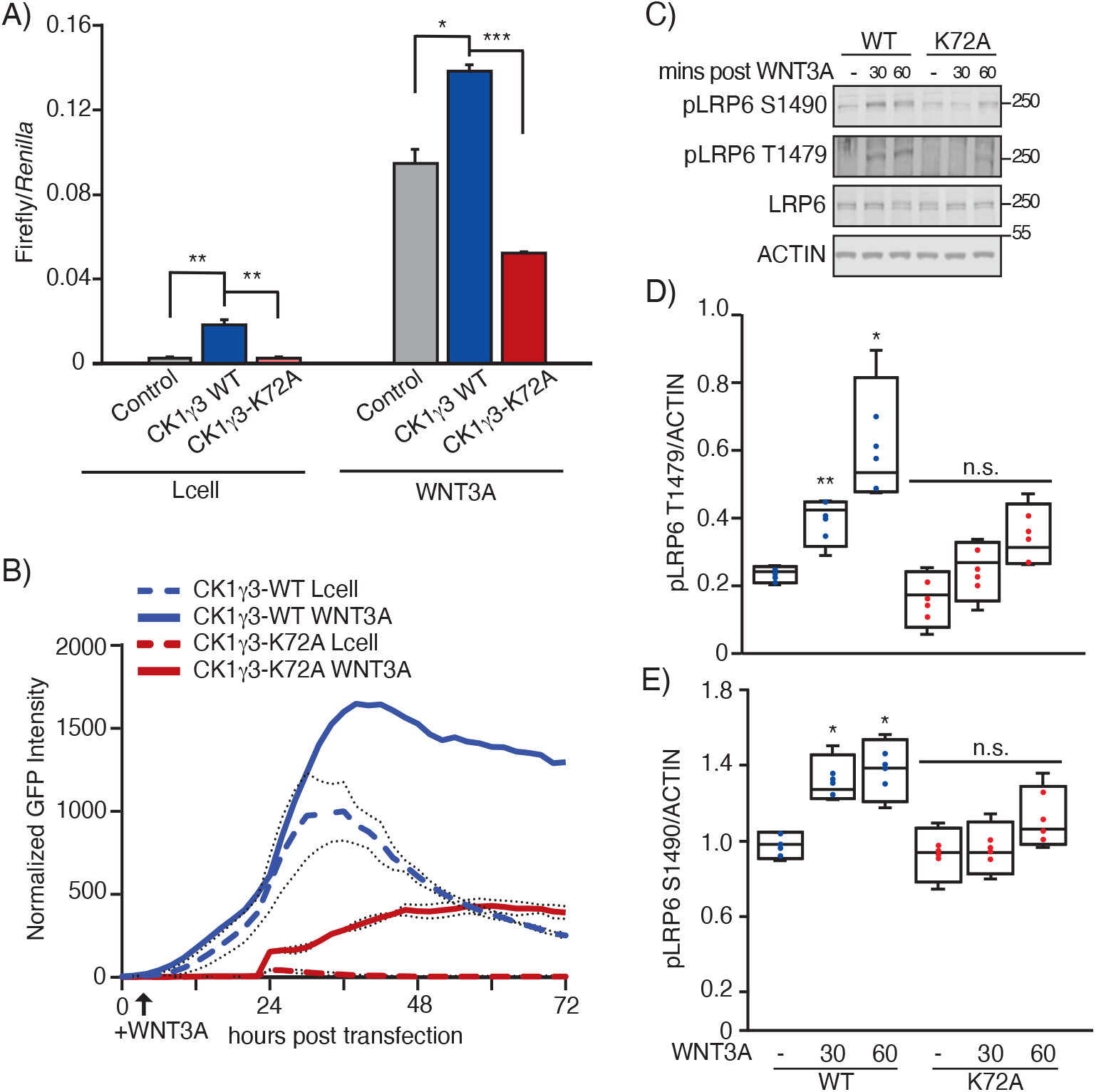
CK1γ3 kinase activity is required for maximum WNT3A activation. **(A)** HEK293T B/R cells were transfected with the indicated constructed for 14 hrs, treated with Lcell or WNT3A CM for 18 hrs, then analyzed by Luciferase assay. **(B)** Live-cell imaging of HEK293T cells stably expressing a BAR-GFP fluorescent reporter transiently transfected with the indicated expression construct, CK1γ3-WT or CK1γ3-K72A. Lcell or WNT3A CM was added at 8 hr, and cells were monitored for an additional 72 hr. Data are averaged across four technical replicates. **(C)** HEK293T cells transfected with either CK1γ3-WT or CK1γ3-K72A for 14 hrs then either treated with WNT3A CM or Lcell CM for 30 or 60 mins. Samples were then analyzed by W. blot. **(D, E)** Box-and-whisker plot quantification of W. blot from panel A for indicated time points, pLRP6 T1479 (panel D) and pLRP6 S1490 (panel E) across 4 replicates, normalized to total LRP6. All statistics are compared to Flag-CK1γ3 Lcell sample. For all panels: *** p<0.0005, ** p<0.005, * p<0.05 and all data are representative of biological triplicates, unless otherwise noted. Error bars represent S.E.

To extend this finding, we tested the effect of CK1*γ* knockdown on BAR reporter activity. First, we confirmed knockdown by qPCR of two non-overlapping siRNAs for CK1*γ*3, as well as pooled siRNAs targeting CK1*γ*1/2/3 or CK1*γ*1/2 (**Fig. 5A**). Transfection of the indicated siRNA into HEK293T cells stably harboring BAR revealed that knockdown of CK1*γ*3 alone or a pool of CK1*γ*1/2 did not significantly inhibit WNT activation. However, knockdown of CK1*γ*1/2/3 suppressed WNT activation by approximately 50% (**Fig. 5B**). Second, we examined *β*-catenin protein stabilization and LRP6 phosphorylation following a WNT3A time course. *β*-catenin protein levels increased in the control cells. CK1*γ*3 knockdown modestly suppressed WNT3A CM induced *β*-catenin stabilization. With simultaneous knockdown of CK1*γ*1/2/3, stabilization of *β*- catenin was significantly impaired as compared to control siRNAs (**Fig. 5C, D**). Finally, knockdown of CK1*γ*3 and CK1*γ*1/2/3 decreased phosphorylation of LRP6 at T1479 and S1490 compared to the control knockdown (**Fig. 5C, E, F**).

**Figure 5.**
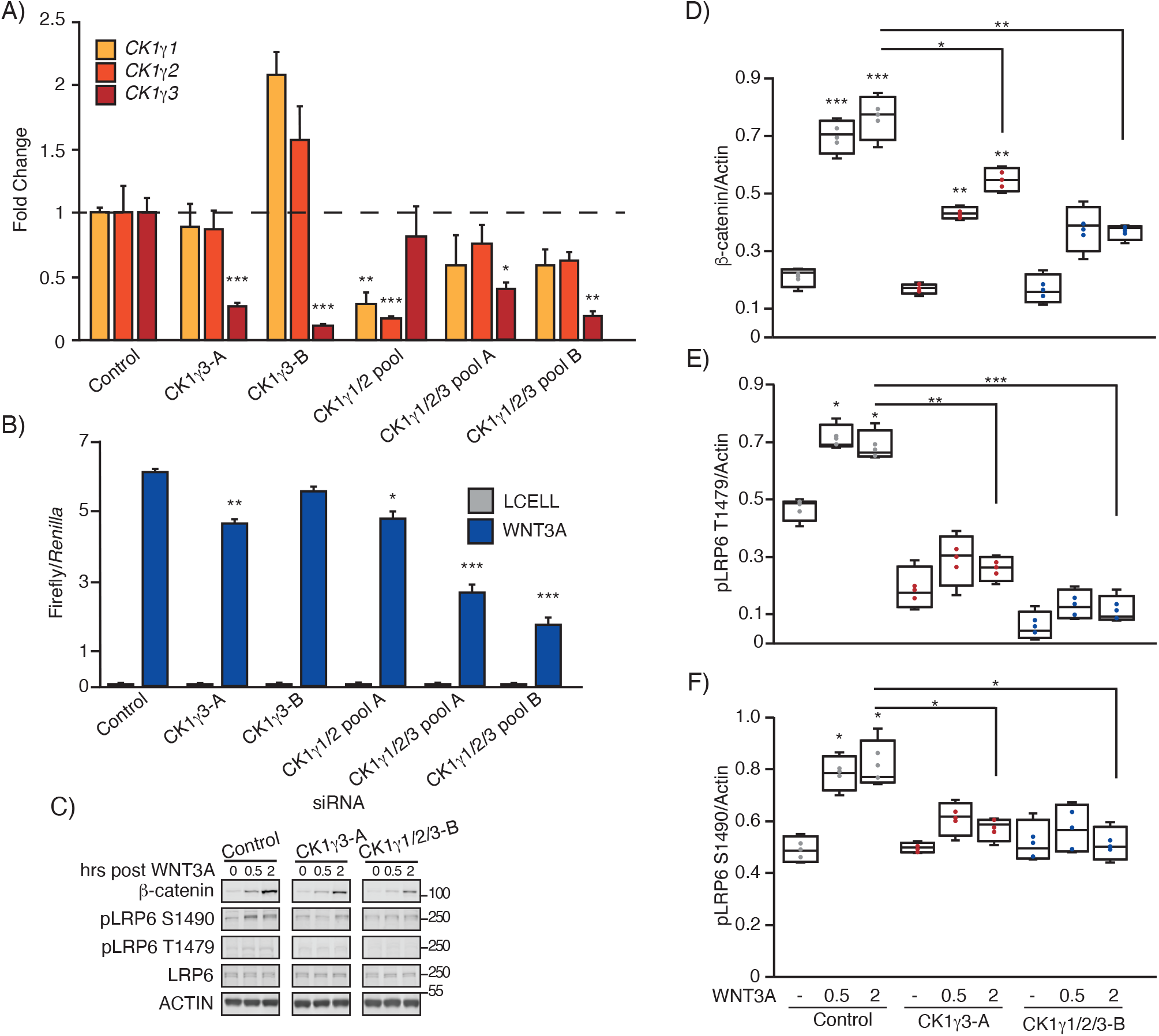
Loss of CK1γ1/2/3 impairs WNT activation. **(A-C)** RKO B/R cells were transfected with indicated siRNA or pooled siRNA for 48 hrs, then split for the following experiments: 24 hrs post-split gene expression was analyzed by RT-qPCR (panel A), 6 hrs post-split cells were treated with WNT3A for 18 hrs then analyzed by Luciferase assay (panel B), 24 hrs post-split cells were exposed to a WNT3A time course, harvested at the indicated time post WNT3A and analyzed by W. blot. For panel A all statistics are compared to siControl expression for the specific gene. For panel B all statistics are compared to siControl WNT3A treated cells. **(D-F)** Box-and-whisker plot quantification of W. blot from panel E, β-catenin (panel D), pLRP6 T1479 (panel E) and pLRP6 S1490 (panel F) across 4 replicates, normalized to ACTIN. Statistics are compared to siControl untreated, unless otherwise stated. For all panels: *** p<0.0005, ** p<0.005, * p<0.05 and all data are representative of biological triplicates. Bars represent average Firefly/*Ren* (RFU) from 3 technical replicates +/- S.E.

### CK1γ3 inhibition impairs activation of WNT signaling

To complement the genetic study of CK1*γ* family within the WNT pathway, we sought CK1*γ* chemical inhibitors. Molecular tool compounds that target the larger CK1 family of kinases have been studied within WNT signaling in the past (58). However, given the multiple positive and negative roles of various CK1 family members in the pathway, and the lack of selectivity for most tool compounds, the results have been inconclusive (59). We leveraged a library of modestly selective kinase inhibitor tool compounds from the University of North Carolina-Structural Genomics Consortium (UNC-SGC), including a 5-substituted indazole with previously characterized inhibition activity for CK1 (60). To further characterize the lead candidate, we sent AKI00000062a (**Fig. 6A, B**) (referred to as AKI throughout) for kinome-wide profiling against 403 WT kinases at DiscoverX at 1 μM. This compound demonstrated modest selectivity, with a calculated S_10_(1 μM) = 0.042, corresponding to 17 kinases with a Percent of Control (PoC) < 10 at 1 μM (**Fig. 6C**). GSK3α and GSK3β were among those kinases potently inhibited, as well as all three CK1*γ* kinases (**Fig. 6D**). To confirm this, we collected the corresponding enzymatic assay activity for all three CK1*γ* family members and found AKI to inhibit all three enzymes with IC_50_ values ≤275 nM (**Fig. 6B**). Finally, a cellular target engagement assay (NanoBRET) was used to measure the potency of enzymatic inhibition for CK1*γ*2 in live cells. We observed modest suppression of CK1*γ*2 in the cell-based system: IC_50_ = 991 nM (**Fig. 6E**).

**Figure 6.**
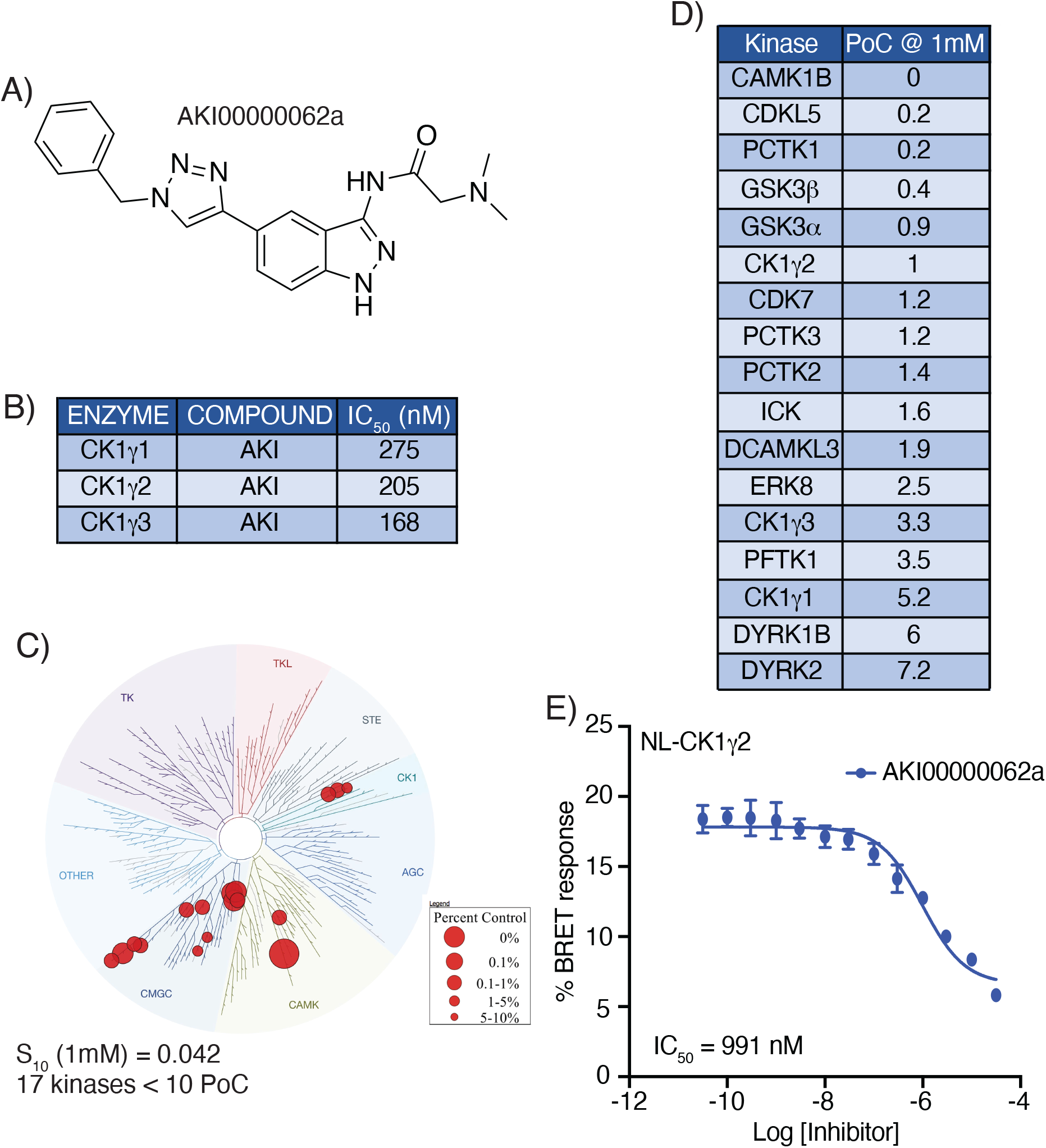
– Summary of data for AKI00000062a. **(A)** Structure of AKI00000062a. **(B)** Enzymatic profiling of AKI00000062a versus CK1γ family kinases. **(C)** Kinome tree representation of kinases with PoC < 10 when AKI00000062a was profiled at 1 μ M versus 403 wild-type kinases. **(D)** Specific kinases (17) with PoC < 10 in kinome-wide profiling. **(E)** Nanoluc-CK1γ2 Nano- BRET curve and corresponding IC50 value for AKI00000062a.

Because the AKI compound also targeted GSK3*β*, we turned to the literature for an alternative chemotype with published potency against the CK1*g* kinase subfamily (58). Amgen released a series of pyridyl pyrrolopyridinones as potent and selective CK1*γ* inhibitors (58). We synthesized one of the published leads (compound 13), furnishing FP1-24-2 with >95% purity (**Fig. 7A**) (referred to as FP1 throughout) (58). We first confirmed the activity of this compound versus all three CK1*γ* enzymes (IC_50_ values <15 nM, **Fig. 7B**). While Ambit KINOMEscan data at 1 μM was included as supplemental information in the original publication, kinome-wide selectivity data were lacking (58). We sent FP1 for kinome profiling against 403 wild-type (WT) kinases at DiscoverX at 1 μM. FP1 demonstrated good selectivity with an S_10_(1 μM) = 0.035 (61), corresponding to 14 kinases with a PoC < 10 at 1 μM (**Fig. 7C**). Among the potently inhibited kinases are all three CK1*γ* family members, CK1*δ*, CK1*ε* and CK1*α* (**Fig. 7D**). NanoBRET was used to test the enzymatic (**Fig. 7B**) and binding (**Fig. 7D**) assay activities for the CK1*γ*2 kinase in cells (62). Gratifyingly, potency was maintained in cells with an IC_50_ of 10.7 nM (**Fig. 7E**).

**Figure 7.**
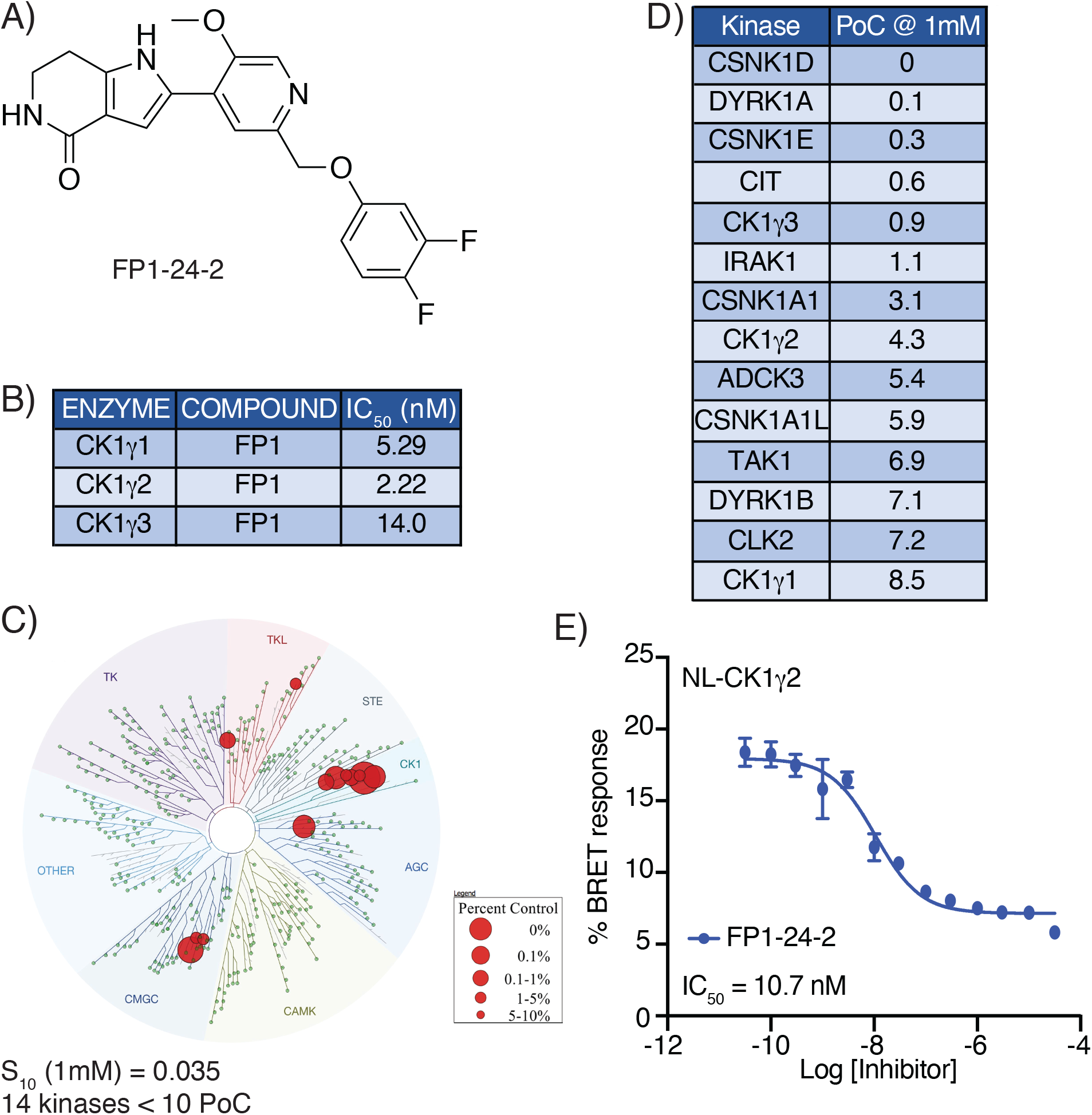
– Summary of data for FP1-24-2. **(A)** Structure of FP1-24-2. **(B)** Enzymatic profiling of FP1-24-2 versus CK1γ family kinases. **(C)** Kinome tree representation of kinases with PoC < 10 when FP1-24-2 was profiled at 1 μM versus 403 wild-type kinases. **(D)** Specific kinases (14) with PoC < 10 in kinome-wide profiling. **(E)** Nanoluc-CK1γ2 Nano- BRET curve and corresponding IC50 value for FP1-24-2.

The kinome-wide profiling of AKI demonstrated potent inhibition of GSK3α and GSK3β; FP1 did not inhibit GSK3α and GSK3β. Using the GSK3β in-cell NanoBRET assay, we found that AKI potently engages GSK3β in cells with an IC_50_ = 7.14 nM (**Fig. 8A**). AKI was significantly more potent on GSK3 kinases than on CK1*γ*2. In contrast, FP1 was inactive up to 10 μM in the GSK3β NanoBRET assay (**Fig. 8B**). As a control, APY69, validated as active in the GSK3 NanoBRET assay by Promega, was included in both the CK1*γ*2 and GSK3β NanoBRET assays. APY69 had an IC_50_ = 0.23 nM in the GSK3β NanoBRET assay, and was inactive in the CK1*γ*2 NanoBRET assay. To confirm that AKI inhibits GSK3*β*, we compared it in a BAR assay with the GSK3*β* inhibitor, CHIR99021 (**Fig. 8C**) (63). Both CHIR99021 and AKI robustly activated the BAR reporter, as compared to DMSO and FP1.

**Figure 8.**
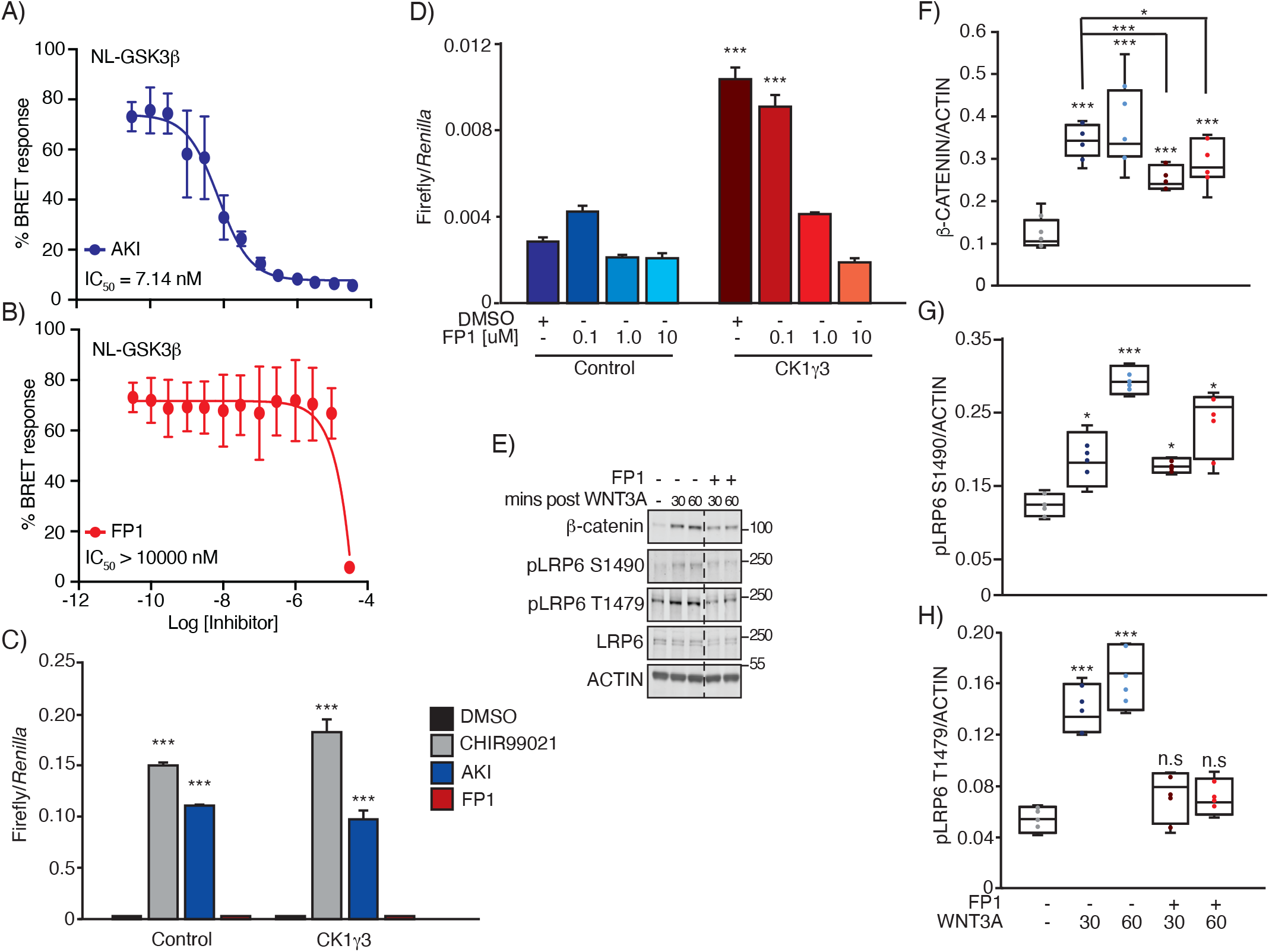
CK1γ inhibition suppresses β-catenin stabilization and LRP6 phosphorylation. **(A)** Nanoluc-GSK3β NanoBRET curve and corresponding IC50 value for AKI. **(B)** Nano- luc-GSK3β NanoBRET curve and corresponding IC50 value for FP1. **(C)** HEK293T B/R cells were transfected with the indicated constructed for 14 hrs, then treated with 10 uM of the indicated compound for 18 hrs and analyzed by luciferase assay. **(D)** HEK293T B/R cells were transfected with the indicated constructed for 14 hrs, treated with DMSO, 0.1, 1.0 or 10 mM of FPI for 18 hrs, and then analyzed by luciferase assay. **(E)** RKO cells were pre-treated with FPI or DMSO for 1 hr, treated with WNT ligand for 30 or 60 mins, then analyzed by W. blot. **(F-H)** Box-and-whisker plot quantification of W. blot from panel E, b-catenin (panel F), pLRP6 S1490 (panel G) and pLRP6 T1479 (panel H) across 4 replicates, normalized to ACTIN. For all panels: *** p<0.0005, ** p<0.005, * p<0.05 and are compared to DMSO control unless other- wise indicated. All data are representative of biological triplicates, unless otherwise noted. Error bars represent S.E. Bars represent average Firefly/*Ren* (RFU) from 3 technical replicates +/- standard error (S.E.).

Finally, we tested whether FP1 impacted CK1*γ*3 activation of WNT signaling in a BAR assay. We overexpressed CK1*γ*3 and treated with increasing doses of FP1, an observed a decrease in BAR activity (**Fig. 8D**). In agreement with this, treatment of cells with FP1 and WNT3A CM resulted in diminished *β*-catenin stabilization and phosphorylation of LRP6 at site S1490, as compared to the DMSO control treated cells (**Fig. 8E, F, G**). Importantly, when cells were treated with FP1 then exposed to WNT3A, LRP6 phosphorylation at T1479 did not increase (**Fig. 8E, H**).

## Discussion

Protein kinases govern information flow though cellular signaling networks, and ultimately impact all of cell biology. Their exceptional druggability and centrality within human disease networks has supported decades of research and therapeutic development (64). However, attention to kinases has not been evenly spread across the protein class. Recently, the NIH Illuminating the Druggable Genome consortium has identified 162 of the most understudied ‘dark’ kinases (33). In this study, we characterize three of these ‘dark’ kinases: CK1*γ*1, CK1*γ*2 and CK1*γ*3. Through unbiased protein-protein proximity network derivation and a functional evaluation across a small focused panel of transcriptional reporters, we connect the CK1*γ* family to the *β*-catenin-dependent WNT signaling pathway. Unexpectedly we found that the CK1*γ*3 family member is uniquely active within *β*-catenin-dependent WNT signaling. All three family members co-complex with both *β*-catenin-dependent and independent WNT machinery. All three family members activate *β*- catenin-dependent transcription in the present of exogenous WNT3A ligand or in LRP6 overexpressing models. CK1*γ*3 appears particularly potent in its activity as it activates *β*-catenin-dependent transcription in the absence of exogenous WNT3A ligand or LRP6 overexpression. We show though siRNA-based silencing and small molecule inhibitors that functional redundancy likely exists within the CK1*γ* family, as CK1*γ*3 suppression alone had minimal impacts on WNT signaling. Beyond connections to the WNT pathway, the proximity networks, functional data and molecular probes offer a deep resource for the kinase community.

Defining proximity networks for this subfamily of understudied kinases provides insight into their cellular functions. Identification of core WNT signaling components, both *β*-catenin dependent and independent, further establishes the CK1*γ* kinase subfamily as WNT signaling components. Many of the identified proximal proteins localize to the plasma membrane, which is consistent with the reported localization of the CK1*γ* family, likely due to palmitoylation of their C-terminus (31). One interesting protein identified as abundant and proximal to all three family members is ZDHHC8, a palmitoyltransferase with strong disease connections to schizophrenia (65). In *Drosophila*, the ZDHHC8 ortholog palmitoylates Scribble, which is a key component of the planar cell polarity pathway and has been shown to negatively regulate WNT/*β*-catenin signaling in human cell lines (66–68). It will be important to determine if ZDHHC8 is responsible for palmitoylation and membrane localization of the CK1*γ* family, and reciprocally if CK1*γ* regulates ZDHHC8 activity or localization. A second interesting and abundant discovery across all three family members is PIK3R3 (phosphoinositide-3-kinase regulatory subunit 3). This regulatory subunit of class 1a PI3K binds phosphorylated tyrosine residues to signal downstream of receptor tyrosine kinases and cytokine receptors (69). PI3K-AKT signal transduction is among the most frequently activated and functionally importance pathways in cancer (70). Whether and how CK1*γ* activity influences PI3K signaling and biology is exciting and warrants testing. Broadly, interrogation of kinase proximity networks like those presented for CK1*γ* offer a powerful resource for kinase substrate prediction and definition. Notably, however, proximity-based networks have caveats that must be considered, including: 1) the approach is exquisitely sensitive to over-expression artifacts, 2) rigorous probabilistic scoring approaches against appropriate controls are needed to illuminate true positives from false positive, 3) as will all of proteomics, absence of identification does not mean absence within the sample but rather below the level of detection; and 4) proximity networks do not identify direct binding relationships. For kinases, motif enrichment queries and integration with phospho-proteomic data can seed experiments for the identification of direct kinase substrates (53).

Our observed physical and functional connections to *β*-catenin-dependent and independent signal transduction is unsurprising. Previous studies demonstrated that CK1*γ*1 activated *β*-catenin- dependent WNT signaling in the presence of exogenous WNT3A ligand, or with co-overexpression of LRP6 or DVL (31, 32). Importantly, our data replicate these results (**Fig. 2D, E**). Also similar to our results, without co-overexpression the authors observed minimal activation of WNT signaling with CK1*γ*1 overexpression (31). However, subsequent studies to further articulate the role of the larger CK1*γ* family in WNT signaling has been lacking until now, including loss of function characterizations. One of the surprises of this work is the family-unique ability of CK1*γ*3 to activate *β*-catenin-dependent transcription in the absence of exogenous WNT3A ligand or co-overexpression of LRP6. Interestingly though, the C59 inhibitor experiment (**Fig. 2F**) establishes that CK1*γ*3 requires an autocrine loop of WNT signaling to impact *β*-catenin.

Loss-of-function studies for the CK1*γ* family within the WNT pathway have not been reported, perhaps due to modest or lacking effect sizes. Functional redundancy within the WNT signaling pathway is well-acknowledged, and indeed our data support redundancy for the CK1*γ* family (71–73). This work and previous studies have shown CK1*δ*/*ε* and CK1*γ* can function to phosphorylate LRP6 (31, 32). In our siRNA-based loss-of-function studies, knock down of all three CK1*γ* subfamily members was required to observe maximal suppression of the WNT pathway (**Fig. 5**). Future studies using CRISPR genetic knockouts are needed, as siRNA-based approaches are incomplete, which is particularly problematic for catalytic proteins like kinases. That said, it is possible that with loss of the full CK1*γ* subfamily, CK1*δ*/*ε* may compensate to phosphorylate LRP6. Beyond genetics, we evaluated two tool chemical inhibitors for the CK1 family. The AKI compound also targeted GSK3*β* which complicates the study of CK1*γ* in WNT signaling (**Fig. 6**). The other compound, FP1, did not target GSK3*β*, but suffered lack of specificity within the CK1 subfamily (**Fig. 7**). FP1 also targets CK1*δ*/*ε*, as well as CK1*α*, which governs WNT signaling through phosphorylating *β*-catenin within the destruction complex (**Fig. 7**). However, FP1 holds value as a research tool when studying proximal events in signalosome formation.

The highly conserved nature of the CK1*γ* subfamily makes the unique function of CK1*γ*3 activation of WNT signaling intriguing and raises the question why CK1*γ*1 and CK1*γ*2 do not activate WNT signaling with the same potency. Structurally, the CK1*γ* subfamily is similar with one major exception – CK1*γ*3 has a 33 amino acid insertion close to the C-terminal tail (**Fig. 1A**). Five *β*-sheets on the c-terminus of the CK1*γ* subfamily proteins create an open crescent moon shape (74). However, in CK1*γ*3 the 33 amino acid insertion creates an additional, short *β*-sheet within an intrinsically disordered region (IDR) (74, 75). This *β*-sheet shifts the C-terminal structure so that the *β*-sheet lies in an opening of the crescent moon shape (74). Additionally, IDRs allow the same peptide sequence to interact differently with different functional outcomes and can have a high number of post-translational modifications (75, 76). The functional implications of the altered structure and the role the IDR plays in CK1*γ*3 have not been evaluated. Studies which delete the 33 amino acid insertion are necessary to fully understand the functional and mechanistic implications.

Illuminating understudied proteins, particularly kinases, is critical to furthering our understanding of several signaling cascades. This work highlights the need to understand the role these understudied kinases play in signaling pathways and disease states. Full characterization of the understudied kinases presents a previously untapped pool of therapeutic targets to treat a variety of human diseases and improve patient outcome.

## Methods and Materials

### Cell lines and tissue culture

All cells were cultured at 37 °C and 5% CO_2_. The following cells were obtained from American Type Culture Collection (ATCC, Manassas, VA): HEK293T/17 (human fetus, cat#CRL-11268), RKO (human, gender N/A, cat# CRL-2577), Lcells (mouse male, cat# CRL-2648), WNT3A expressing Lcells (mouse male, cat# CRL-2648). All cells were grown in DMEM with 10% FBS. Each cell line was tested for mycoplasma contamination and passaged no more than 20 passages from the original ATCC stock.

### WNT3A conditioned media

Conditioned media was collected as described by ATCC. Briefly, WNT3A and control Lcells were grown to 100% confluency before fresh media was added conditioned for 48 hrs, and then collected.

### Lysis buffers

RIPA lysis buffer contains 10% Glycerol, 50mM Tris-HCl pH 7.4, 150mM sodium chloride (NaCl), 2mM Ethylenediaminetetraacetic acid (EDTA), 0.1% SDS, 1% Nonidet P-40 (NP-40), 0.2% Sodium Deoxycholate. TAP lysis buffer contains 10% Glycerol, 50 mM HEPES pH 8, 150 mM NaCl, 2 mM EDTA, 0.1% NP-40. Triton lysis buffer contains 10% glycerol, 50mM Tris-HCl pH 7.4, 150mM NaCl, 2mM EDTA, and 1% TritonX1000.

### Cloning

mT constructs (UBC-mT-V5-CONSTRUCT) were generated by LR multisite recombination (Life Technologies, Carlsbad, CA, cat# 11791019; comes with LR Clonase ® II enzyme mix) utilizing the PEL system (77). CK1*γ*3 kinase dead mutants were generated using PCR mutagenesis and cloned into pDONR223 digested with BsrGI using Gibson Assembly Master Mix (New England Biolabs (NEB), Ipswich, MA, cat#E2611).

### CK1γ3 K72A (AAG to GCG)

CK1*γ*3 gib A-F: 5’- CTTTTTTATAATGCCAACTTTGTACAAAAAAGTTGGCATGGAGAACAAGAAGAAGG ACAAGG

CK1*γ*3 K72A gib A-R: 5’- GCTCCAGCGCGATGGCCAC

CK1*γ*3 K72A gib B-F: 5’-GTGGCCATCGCGCTGGAGC

CK1*γ*3 gib B-R: 5’-

TTTCTTATAATGCCAACTTTGTACAAGAAAGTTGGACTACTTGTGTCTCTGGATTGTC TTCC

### Immunoblotting

Standard W. blotting techniques were utilized and performed as previously described (27, 52). Briefly, cells were lysed in RIPA lysis buffer, flash frozen on dry ice, then high speed cleared for 10 mins at max speed. Protein concentration was determined by a BCA assay following kit specifications. Samples were balanced and NuPAGE 4x SDS loading buffer (ThermoFisher, Waltham, MA, cat# NP0007) containing DTT was added to each sample and heated for 10 mins at 70 °C. Samples were run on a NuPAGE 4-12% Bis-Tris Protein Gel (ThermoFisher, cat# NP0321), then transferred to a nitrocellulose membrane. The membrane was then blocked and incubated overnight with primary antibody at 4 °C with rotation, washed with TBST, incubated with secondary antibody, washed with TBST and imaged. Images were taken using a LiCOR Odyssey CLx imager and quantified with LiCOR software (LiCOR Biosciences, Lincoln, NE). All antibodies were used at a concentration of 1:1000, with the exception of loading controls, which were used at 1:10,000. All primary antibodies utilized are as follows: *β*- catenin (BD Biosciences, Franklin Lakes, NJ, cat# 163510), LRP6 (Cell Signaling Technology, Danvers, MA, cat# 3395), phospho LRP6 S1490 (Cell Signaling Technology, cat# 2568), phospho LRP6 T1479 (Life Technologies, cat# PA564736), AXIN1 (Cell Signaling Technology, cat# 3323), GSK3*β* (BD Biosciences, cat# 610201), CK1*γ*1 (Abcam, Cambridge, United Kingdom, cat# ab190260), CK1*γ*2 (Abcam cat# ab64829), CK1*γ*3 (Abcam cat# ab116310), DVL3 (Santa Cruz Biotechnology, Dallas, TX, cat# sc-8027), CELSR2 (Cell Signaling Technology cat# 47061), V5 (ThermoFisher Scientific cat# R960-25), *β*-ACTIN (Santa Cruz Biotechnology cat# sc-47778). Secondary antibodies were used at a concentration of 1:20,000 and were as follows: IRDye 800CW Goat anti Mouse IgG (LI-COR Biosciences, cat# 925-32210), IRDye 680LT Goat anti Mouse IgG (LI-COR Biosciences, cat# 925-68020), IRDye800CW Goat anti Rabbit IgG (LI-COR Biosciences, cat# 925-32211, IRDye 680LT Goat anti Rabbit IgG (LI-COR Biosciences, cat# 925- 68021), IRDye 680LT Streptavidin (LI-COR Biosciences, cat# 926-68031).

### Transcriptional reporter assays

All luciferase reporter assays were performed as previously described (27, 52). Briefly, cell lines stably expressing the BAR-Firefly luciferase reporter and TK-Ren luciferase were used and transfected with either RNAiMAX (Life technologies, cat# 13778150) for 60 hrs in loss of function experiments and Lipofectamine 2000 (Life technologies, cat# 11668019) for 36 hrs in gain of function experiments. In both loss and gain of function experiments, treatments (WNT CM or drug) were added 18 hrs before harvesting cells. Conditions were plated in triplicate, and *Renilla* normalized values were averaged across triplicates to yield the data presented and standard error. Each assay was repeated in biological triplicate, unless otherwise stated. Firefly luciferase and the *Renilla* control were detected using the Promega Dual-Luciferase Reporter Assay System per the manufacturer’s protocol (Promega, Madison, WI, cat# E1960). Plates were read on the Victor Nivo plate reader (Perkin Elmer, Waltham, MA).

### siRNA knockdown

All siRNAs were obtained from Life Technologies (ThermoFisher, Waltham, MA). Cell lines were used and transfected with RNAiMAX (Life technologies, cat# 13778150) and for 72 hrs before W. blot analysis.

### Real-Time Quantitative PCR

Total RNA was extracted from RKO and HT1080 cells using Invitrogen PureLink RNA mini kit (Life Technologies, Cat#12183025) following the manufacture’s manual. The AXIN2 and NKD1 primers are described in (Walker et al 2015). All other primers were designed using the National Center for Biotechnology Information’s (NCBI) Primer-BLAST platform. Primer sequences are as follows: CK1*γ*1 Forward 5’ CCTCATTTGCGCCTTGCAG 3’ Reverse 5’ CTCCGGGAGATGAAAAACCA 3’; CK1*γ*2 Forward 5’ GGCGGAGCCAAGCTGTGA 3’ Reverse 5’ GCGCTGCTCTTCTCAGTGG 3’; CK1*γ*3 Forward 5’ TTGCAGCAGACAGACATGGT3’ Reverse 5’ GGCTGTGATGGGTGCATTTG 3’; and RPL13a Forward 5’ CATAGGAAGCTGGGAGCAAG 3’ Reverse 5’ GCCCTCCAATCAGTCTTCTG 3’. Before reverse transcription, RNA was quantified using a Nanodrop One (Thermo, Waltham, MA), and 1 μg of RNA was used to generate cDNA with the iSCRIPT Clear kit (Bio-Rad, Hercules, CA, cat# 170-8891). For the qRT-PCR, PowerUP SYBR Green (Thermo, cat# A25741) was used, and data were analyzed on a QuantStudio 5 Real Time PCR machine (Applied Biosystems, Foster City, CA). ΔCT values were normalized to housekeeping gene RPL13a.

### Affinity pulldowns

Cells were incubated for indicated time with 50mM biotin, washed 3x in cold PBS, then lysed in RIPA lysis buffer, flash frozen on dry ice, then thawed and incubated on ice for 30 mins. Lysates were then sonicated for 20 seconds in 5 second pulses. Samples were then high speed cleared and protein concentration was determined by BCA. 50uL of bead slurry per sample was washed 5x in RIPA lysis buffer, then incubated with lysate at 4 °C with rotation for either 1 hr for W. blot analysis or overnight for mass spectrometry analysis. Beads were then washed 5x with RIPA lysis buffer. For W. blot analysis samples were then eluted in a 1:1:1 mixture of 1 M DTT, 4x SDS loading buffer and RIPA lysis buffer and heated at 70 °C for 10 mins.

### Affinity purification and mass spectrometry

Following the affinity pulldown (described above), beads were washed 2x with RIPA, 2x with TAP lysis buffer, 3x with 50mM ammonium biocarbonate (ABC) buffer. 1mg of RapiGest SF surfactant (Waters Corporation, Milford, MA, cat# 186001861) was resuspended in 200uL of 50mM to give a 0.5% RapiGest solution. Streptavidin beads were resuspended in 100uL of 0.5% RapiGest and vortexed. 1M Dithiothreitol (DTT) in ABC was added to the sample for a final concentration of 5mM and samples were heated at 60°C for 30 mins. After allowing samples to cool to room temperature, chloroacetamide was added to each sample for a final concentration of 15mM and incubated in the dark for 20 minutes at room temperature. The samples were then centrifuged at 400 x g for 2 mins at room temperature to remove the supernatant containing protein from the streptavidin beads. The supernatant was transferred to a new tube and 2.5 μg of mass spectrometry grade trypsin was added to each sample. The samples were incubated at 37°C overnight, then Hydrochloric acid (HCl) was added to the sample at a final concentration of 250mM and incubated for 45 mins at 37°C to deactivate the trypsin. Pierce C18 spin columns (ThermoFisher, cat#89870) were placed in a 2 mL centrifuge tube, the column was activated by adding 200 μL of 100% acetonitrile (ACN) and centrifuged at 3000 xg for 2 mins. Columns were equilibrated by adding 200 μL of 0.5% Trifluoroacetic acid (TFA) and centrifuged at 3000 xg for 2 mins. This step was then repeated. The sample was resuspended in 200 μL of 0.5% TFA, then loaded onto the column and centrifuged for 5 mins at 1000 xg. The sample was reloaded onto the column and centrifuged a second time for 5 mins at 1000 xg. The column was washed twice with 200 μL of 5% ACN/0.5% TFA and centrifuged for 2 mins at 3000 xg. The samples were eluted by adding 50 μL 70% ACN and centrifuged for 5 mins at 1000 xg. A second elution step was performed by adding 50 μL 70% ACN and centrifuged for 2 mins at 3000 xg. Following a C18 clean up, an ethyl acetate clean-up was performed. The sample was resuspended in 100 μL of 0.1% TFA, then 1 mL of water saturated ethyl acetate was added to each sample, vortexed and centrifuged at max speed for 5 mins. The upper layer of ethyl acetate was removed. This process was repeated for a total of 3 times. Samples were then dried down in the speed vac and resuspended in 25 μL of 98/2 Buffer A (Water + 0.1% FA)/Buffer B (ACN + 0.1% FA).

### Mass spectrometry data acquisition

Trypsinized peptides were separated via reverse-phase nano-HPLC using an RSLCnano Ultimate 3000 (Thermo Fisher Scientific). The mobile phase consisted of water + 0.1% formic acid as buffer A and acetonitrile + 0.1% formic acid as buffer B. Peptides were loaded onto a µPACᵀᴹ Trapping column (PharmaFluidics) and separated on a 200 cm µPACᵀᴹ column (PharmaFluidics) operated at 30°C using a 100 min gradient from 2% to 25% buffer B, followed by a 20 min gradient from 25% to 35% buffer B, flowing at 300 nL/min. Mass spectrometry analysis was performed on an Orbitrap Eclipse (Thermo Fisher Scientific) operated in data-dependent acquisition mode. MS1 scans were acquired in the Orbitrap at 120k resolution, with a 250% normalized automated gain control (AGC) target, auto max injection time, and a 375- 1500 m/z scan range. Both the linear ion trap and the Orbitrap were used for MS2 scans. MS2 targets with a charge of +2 or +3, and >90% precursor fit at either a 0.8 m/z or 0.4 m/z-wide isolation window were fragmented with by collision induced dissociation (CID) at 35% collision energy and scanned in the linear ion trap at the widest window width that passed the thresholds. All remaining MS2 targets with charge from +2 to +6, >5e4 intensity, and >10% precursor fit at isolation widths of 1.6 m/z, 0.8 m/z, or 0.4 m/z wide were fragmented with higher-energy collision dissociation at 30% collision energy and scanned in the orbitrap at 15k resolution at the widest window width that passed the thresholds. MS2 AGC targets and maximum injection times were set to standard and auto for their respective analyzers. Dynamic exclusion was set to 60 seconds. Acquisition was performed with a 2.7 second cycle time.

### Mass spectrometry data processing

Raw MS data files were processed by MaxQuant (version 1.6.17.0) (78) using the UniProtKB/SwissProt human canonical sequence database (downloaded Aug. 2019) (79). To facilitate comparison to another 44 bait experiments performed within our laboratory, the files were processed simultaneously with these raw files. The following parameters were used: specific tryptic digestion with up to two missed cleavages, fixed carbamidomethyl modification, variable modifications for protein N-terminal acetylation, methionine oxidation, match between runs, and label-free quantification. Only unique peptides were used for protein quantification due to the high tryptic peptide overlap between CK1*γ*1, CK1*γ*2, and CK1*γ*3. Baits were assigned fractions numbers >1 away from each other to enable match-between-runs within replicates, but not across different baits. Prey proteins were filtered for high-confidence physical interactions and proximal proteins using SAINTexpress (v3.6.3) (45, 46) with the following thresholds: ≥2 unique peptides, AvgP >0.7, and BFDR *≤*0.05. Interactions were further refined by selecting the top 10% of interactions as scored by the CompPASS WD score (https://github.com/dnusinow/cRomppass), validated WNT-related proteins from the literature,and hits from published siRNA and CRISPR screens for WNT pathway regulation (47–51). The criteria for screen hits were as follows: ≤0.01 FDR for Lebensohn et al., ≥2-fold change for Bichele et al., and ≥2-fold change for at least 2 siRNAs in Major et al. Gene Ontology annotations were extracted with the org.Hs.eg.db R package, and overrepresentation analysis was performed with ClusterProfiler using default parameters (80, 81).

### Source of compounds

FP1-24-2 was prepared as previously described (58). Details are provided in the next section. AKI00000062a was donated to SGC-UNC by AbbVie.

### Synthesis of 2-(2-((3,4-Difluorophenoxy)methyl)-5-methoxypyridin-4-yl)-1,5,6,7-tetrahydro- 4*H*-pyrrolo[3,2-*c*]pyridin-4-one (FP1-24-2)

To a flask was added 2-bromo-1-(2-((3,4- difluorophenoxy)methyl)-5-methoxypyridin-4-yl)ethan-1-one (26.3 mg, 1.0 equiv., 70.7 µmol), piperidine-2,4-dione (9.59 mg, 1.2 equiv, 84.8 µmol), and ammonium acetate (21.8 mg, 4.0 equiv, 283 µmol) in ethanol (283 µL, 0.25 molar) and the reaction stirred at rt for 16 hrs. The reaction was concentrated *in vacuo* and purified by preparative high performance liquid chromatography (10-100% MeOH in H_2_O (+ 0.05%TFA) to yield the desired product 2-(2-((3,4- difluorophenoxy)methyl)-5-methoxypyridin-4-yl)-1,5,6,7-tetrahydro-4*H*-pyrrolo[3,2-*c*]pyridin- 4-one as an off-white solid (1.9 mg, 7%).

^1^H NMR (400 MHz, MeOD-*d*_4_): δ 8.30 (s, 1H), 7.77 (s, 1H), 7.22 - 7.13 (m, 2H), 6.99 (ddd, *J* = 3.0, 6.7, 12.4 Hz, 1H), 6.85 – 6.80 (m, 1H), 5.08 (s, 2H), 4.09 (s, 3H), 3.57 (t, *J* = 7.0 Hz, 2H), 2.97 (t, *J* = 7.0 Hz, 2H). ^13^C NMR (101 MHz, MeOD-*d*_4_): δ 169.0, 152.1, 150.1, 140.8, 134.4, 130.3, 127.7, 119.3, 118.48, 118.46, 118.29, 118.28, 115.5, 111.74, 111.70, 111.68, 111.64, 110.5, 105.7, 105.5, 72.0, 57.0, 41.7, 23.0. ^19^F NMR (376 MHz, MeOD-*d*_4_): δ -138.23 – -138.37 (m), - 150.69 (dddd, *J* = 3.3, 6.6, 10.2, 20.8 Hz). LCMS: expected mass for [M+H]^+^ (C_20_H_18_F_2_N_3_O_3_), 386.12 *m/z*; found, 386 *m/z*. LCMS and ^1^H NMR: >95%.

### CK1γ family enzymatic assays

Eurofins kinase enzymatic radiometric assays were carried out at the K_m_ = ATP in dose–response (9-pt curve in duplicate) for each CK1*γ* kinase listed in **Fig.s 6B and 7B**. Details related to the substrate used, protein constructs, controls, and assay protocol for each kinase assay are included on the Eurofins website: https://www.eurofinsdiscoveryservices.com.

### Kinome-wide profiling

The *scan*MAX assay platform was used to assess the selectivity of FP1- 24-2 and AKI00000062a at 1 µM at Eurofins DiscoverX Corporation. As described previously, this commercial assay platform screens against 403 WT human kinases in binding assays and provides PoC values (82). All WT kinases that demonstrated PoC values < 10 are plotted on the respective kinome trees in **Fig.s 6C**, **7C**. These same WT kinases with PoC value < 10 are tabulated in **Fig.s 6D, 7D**.

### NanoBRET assays

Human embryonic kidney (HEK293T) cells (hypotriploid, female, fetal) were purchased from ATCC. These cells were grown in Dulbecco’s Modified Eagle’s medium (DMEM, Gibco) supplemented with 10% (v/v) fetal bovine serum (FBS, Corning). They were incubated in 5% CO_2_ at 37°C and passaged every 72 hours with trypsin and not allowed to reach confluency. Constructs for NanoBRET measurements of CK1*γ*2 (NLuc-CK1*γ*2) and GSK3β (NLuc-GSK3β) were kindly provided by Promega. NanoBRET assays were carried out as described previously (83). Preferred NLuc orientations, both N-terminal in this case, are indicated in parentheses after each construct. Assays were carried out in dose–response as described by the manufacturer using 0.5 μM of tracer K-10 for CK1*γ*2 and 0.063 μM of tracer K-8 for GSK3β. Two biological replicates were executed with two technical replicates to produce the graphs with error bars shown in **Fig.s 6E, 7E, 8A, 8B**. Tracer titration curves for these kinases that motivated our tracer selection and working concentration can be found on the Promega website.

### Supporting information

This article contains supporting information.

## Acknowledgements

Constructs for NanoBRET measurements of CK1*γ*2 and GSK3β were kindly provided by Promega. We used the TREE*spot* kinase interaction mapping software to prepare the kinome trees in **Fig.s 6C and 7C**: http://treespot.discoverx.com.

## Funding and additional information

MJA was funded by HHMI Gilliam Fellowship and the NIH NCI Predoctoral to Postdoctoral Fellow Transition Award (F99/K00 Fellowship (5F99CA245724-02)). This work was supported in part by a grant from the National Institutes of Health Illuminating the Druggable Genome Program [U24DK116204]. The Structural Genomics Consortium is a registered charity (number 1097737) that receives funds from AbbVie, Bayer Pharma AG, Boehringer Ingelheim, Canada Foundation for Innovation, Eshelman Institute for Innovation, Genome Canada, Genentech, Innovative Medicines Initiative (EU/EFPIA) [ULTRA-DD grant no. 115766], Janssen, Merck KGaA Darmstadt Germany, MSD, Novartis Pharma AG, Ontario Ministry of Economic Development and Innovation, Pfizer, São Paulo Research Foundation-FAPESP, Takeda, and Wellcome [106169/ZZ14/Z]. Research reported in this publication was supported in part by the NC Biotechnology Center Institutional Support Grant 2018-IDG-1030, NIH 1U24DK116204 and NIH U54AG065187.

## Supplemental Information

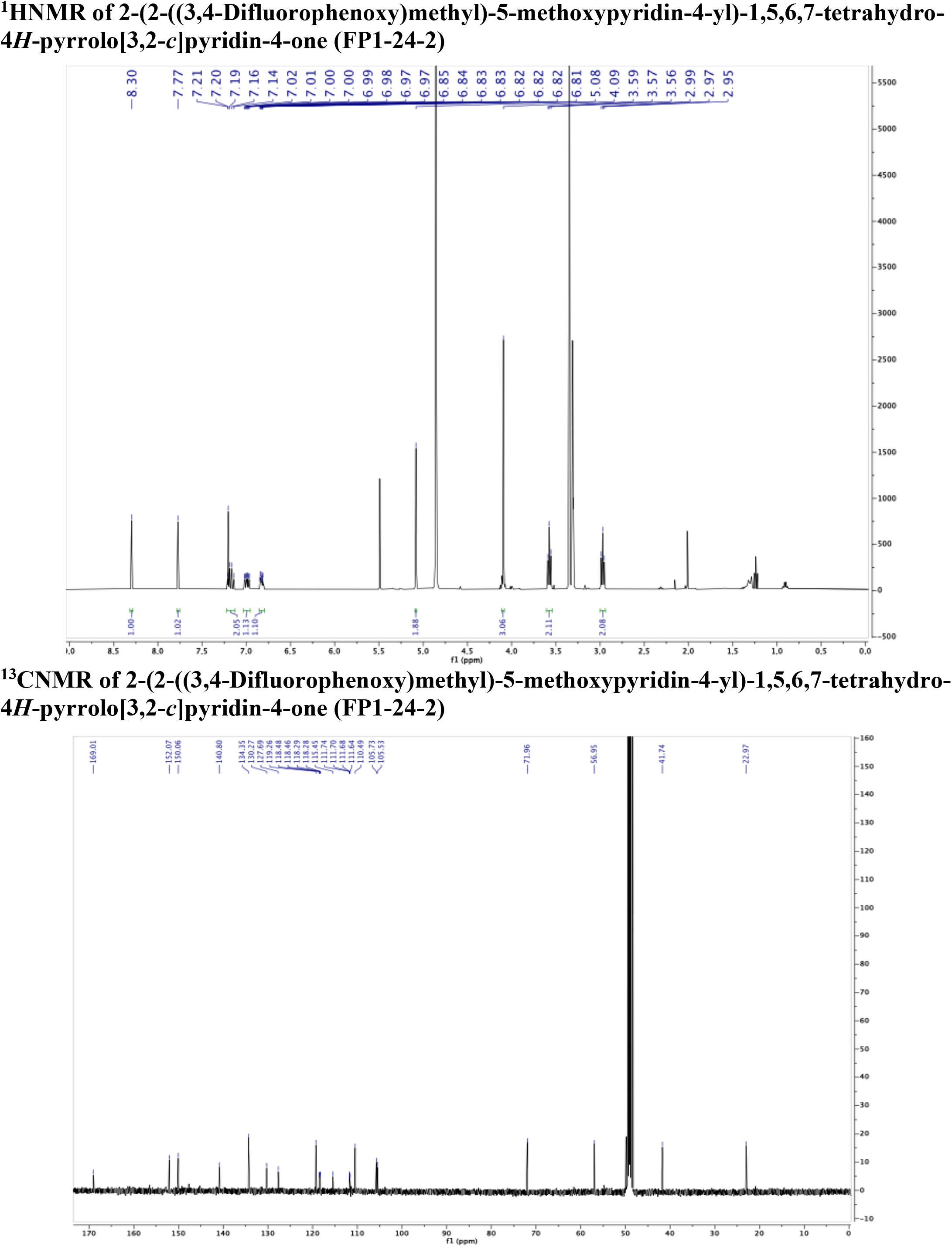

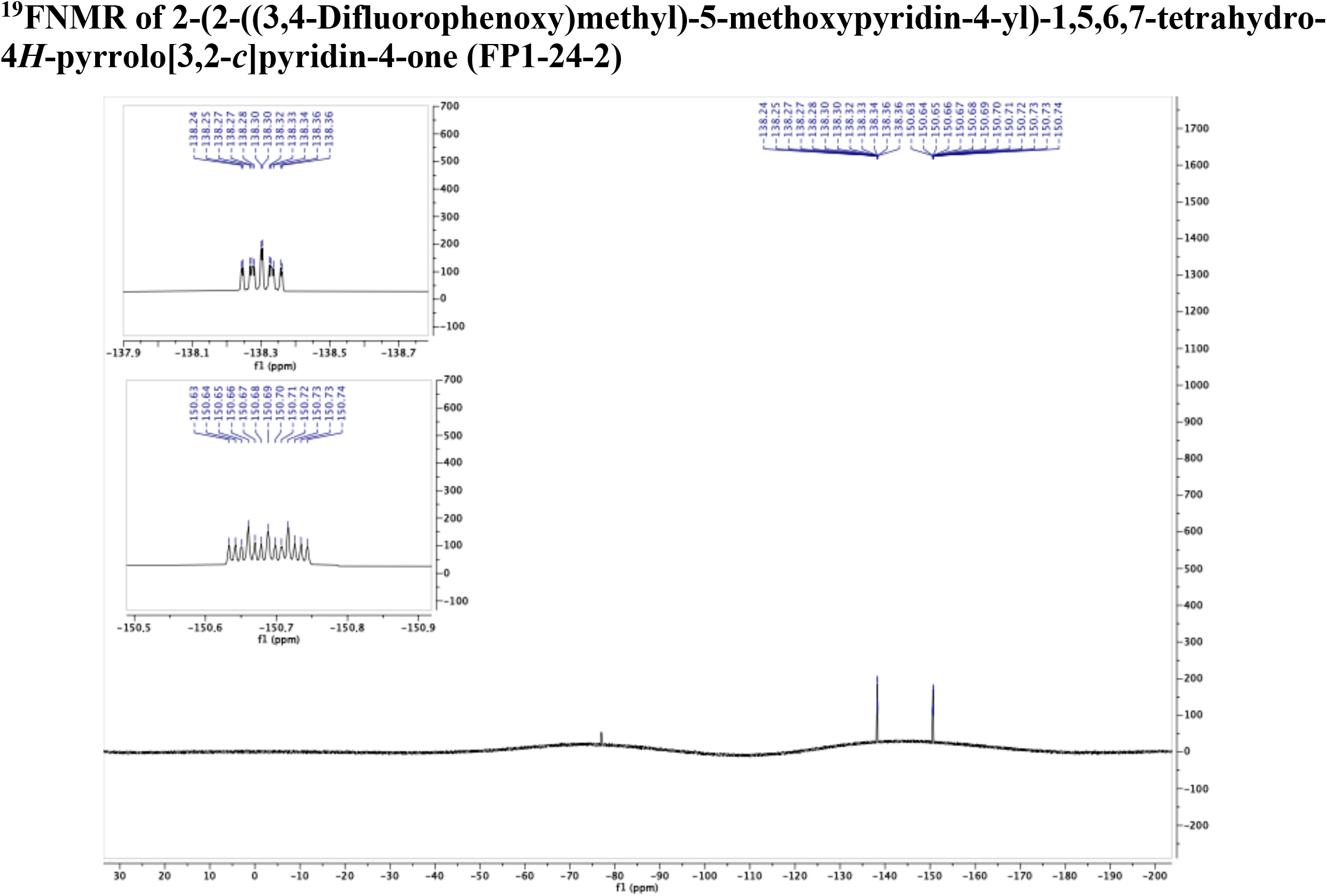

